# H_2_O_2_ mediated MAPK-mTOR crosstalk determines the XIAP and FLIP degradation pathways and development of resistance to Imatinib

**DOI:** 10.1101/2024.04.28.591571

**Authors:** Rajdeep Roy, Tamalika Paul, Samraj Sinha, Nabendu Biswas

## Abstract

Abnormal or aggregated proteins degradation is very crucial to protect the cell against proteotoxic stress. Cellular proteins are usually degraded in a ubiquitin-dependent manner by proteasomes and macro-autophagy/autophagy. Both processes use ubiquitin, binding of which target them to proteasomes via ubiquitin-like domains or to phagophores (the precursors to autophagosomes) via Atg8/LC3 binding motifs. Since both pathways use ubiquitin, the question that arises is how degradative pathway choice is achieved. Here in this paper, we have shown that mTOR pathway acts as major player in deciding which degradation pathway to choose. Here, we used Imatinib resistant (K562R) and Imatinib-sensitive (K562S) K562 cell as our previous study showed that Ubiquitin-proteasome pathway is selected for degradation of two proteins XIAP and FLIP in Imatinib-resistant K562 cells. H_2_O_2_ treatment in K562R cells degrades XIAP and FLIP via proteasomal pathway whereas H_2_O_2_ degrades them via lysosomal pathway in K562S cells. We found H_2_O_2_ mediated mTOR activation suppressed autophagy pathway in K562R cells and protein takes proteasomal degradation rout. However, proteins get degraded via autophagy when mTOR remains suppressed in K562S cells. We also found that H_2_O_2_-mediated activation of activation of ERK in turn activated mTOR in K562R cells because ROS inhibited ERK phosphorylation and thus mTOR remained inhibited in K562S cells. Moreover, we found that endogenous H_2_O_2_ levels are way higher in K562R cells than in K562S cells. Finally, and most interestingly, our data clearly suggested that activation of mTOR is responsible for developing resistance to Imatinib in K562R cells. Thus, this paper delineates a very crucial role of H_2_O_2_ and mTOR in both the selection of protein degradation pathway and development of drug resistant.

## 1. Introduction

Every cell requires a stable proteome for its survival, for which precise regulations of protein synthesis and degradation of abnormal and unwanted proteins are equally important. Degradation of aggregated and abnormal non-functional proteins is vital to shield the cell against the proteotoxic pressure. It is evident that at a given moment nearly one-third of newly synthesized proteins in a cell are degraded. In such a high protein turnover, cell must have quick adaptive system to maintain cellular integrity and endurance[1].

The ubiquitin–proteasome system (UPS) and autophagy are the two major protein degradation pathways in eukaryotes. Upon environmental stimuli both pathways orchestrate several cellular processes. However, ubiquitination of target proteins is used as a ‘death’ signal by both processes. Interestingly, current findings exposed an unswerving functional link between these two pathways[2]. Cell has very intricate signaling network to control which pathways should get activated in any given cellular conditions.

Preeminent levels of 26S proteasome, and remarkably high levels of proteasome activity, have been perceived in numerous different types of cancer. Many recent studies showed that high proteasome activity seems to be important for cancer cell survival, as it possibly supports in protection against apoptosis pathways and eliminate the damaged proteins. Therefore, inhibition of the proteasome is an area of interest in cancer research. In a variety of cancers proteasome inhibition has been shown to persuade apoptosis[3,4].

On the other hand, Autophagy is an exceedingly preserved cellular process in which aggregated or misfolded proteins, intracellular pathogens, and damaged organelles, are degraded and re-used. It has been observed autophagy is dysregulated in different pathological conditions, like neurological disorders, cancer, infection, and aging. However, autophagy and apoptosis epitomize two self-regulatory mechanisms by which cells respond to diverse types of stresses and death stimuli and maintain homeostasis[5–7]. The knowledge is very limited concerning the crosstalk between these two pathways in the primary stages of cancer or under any stress conditions.

Besides being a stress molecule, Hydrogen peroxide (H_2_O_2_) has grasped an appreciable benchmark in cellular signaling in recent years. H_2_O_2_, plays a critical role in controlling many signaling pathways by oxidative modulation of redox sensitive proteins denominated as redox switches[8]. Proteins containing cysteine residues and metal centers are two main redox sensitive moieties. Many proteins including kinases, protein phosphatases and many transcription factors harboring redox switches are important players in the regulation of various biochemical pathways[9]. ROS have a direct impact on several signaling pathways by interacting with many proteins, involved in the control of apoptosis and cell proliferation. As we previously discussed ROS amount/concentration is a chief factor in the growth, development, and cancer cells stemness. Additionally, cancer cells uphold a subtle balance between ROS and antioxidants to endorse pathogenesis and clinical challenges via targeting a series of signaling pathways in cancer. Current findings also demand the therapeutic reputation of ROS for the improved clinical consequences in cancer patients as they persuade apoptosis and autophagy[10,11].

Recent work by our group depicts that hydroxychavicol (a polyphenol from piper betel leaf) were able to induce apoptosis in TRAIL resistant CML cells (K562) by downregulating FLICE Like Inhibitor of Apoptosis Protein (FLIP) and X-Linked Inhibitor of Apoptotic Proteins (XIAP), two of the major IAP proteins (inhibitor of apoptotic proteins) under the influence of ROS (Reactive Oxygen Species)[12]. Another study from our lab showed how ROS treatment induce MAPK pathway component ERK and AKT which involve in XIAP downregulation and in other hand increased expression of ITCH (E3 ubiquitin ligase) involved in FLIP down regulation via proteasomal degradation pathway in drug resistant CML cells[13]. Here in this report, we tried to find out how ROS affects the protein homeostasis by manipulation of its degradation pathways. We emphasized the degradation of XIAP and FLIP under H_2_O_2_ stress and used two Chronic Myeloid Leukemia (CML) cell lines - Imatinib resistant K562 (K562R) and Imatinib sensitive k562 (K562S), to delineate the differential signaling mechanisms behind these control of degradation pathways.

## 2. Materials

Antibodies like c-FLIP, XIAP, PI3K, pPI3K, Akt, pAkt, JNK, pJNK, ERK, ITCH, anti-Rabbit IgG & anti-mouse HRP-linked antibody, Control siRNA, SAPK/JNK siRNA and ERK1/2 siRNA were obtained from Cell Signaling Technology (Denver, Massachusetts, USA). Propidium iodide was bought from Sigma-Aldrich. SureBeads™ Protein G Magnetic Beads was bought from BIO RAD (USA). FITC Annexin V was taken from BD Pharmingen™. JNK inhibitor SP600125, p38 inhibitor SB203580, ERK inhibitor PD98059 were obtained from Abcam (Cambridge, MA, USA). Fetal Bovine Serum (FBS) Standard (origin South America) was procured from Gibco®.Life Technologies, USA. iScript™ Reverse Transcription Supermix for RT-qPCR and SsoFast™ Evagreen® Supermix were purchased from BIO RAD (USA).

## 3. Methods

### Cell Line and cell culture

K562 CML cell lines, both imatinib sensitive and resistant were maintained in RPMI medium which was complemented with 1% penicillin-streptomycin and 0.1% gentamycin and 10% FBS. In case of resistant cells 1.5 μg/ml Imatinib was added for maintaining the resistance against the drug. All cells were cultured at 37° C humidified condition with continuous supply of 5 % carbon dioxide (CO2).

### Western blot analysis

After the experiment is over specified in the respective figure legends, all cells were washed with 1X filtered PBS twice before lysing it with 1X RIPA (CST, Cat no 9806S), supplemented with the protease inhibitor. Cells were subjected for sonication for 15 seconds then cell lysate were collected using high speed centrifugation 15000 rpm for 16 minutes. The concentration of proteins was measured using by Lowry (Folin-Ciocalteau) method. Proteins were subjected to SDS-PAGE by boiling it with Laemmli buffer at 95 ◦C for 3-5 min and. PVDF membrane was used to transfer the proteins for following steps and 5% non-fat dried milk diluted in TBST (2 hours) were used to block the membrane and incubated in respective antibodies overnight in a moist chamber. For detection HRP linked corresponding secondary antibodies were used. Bands were developed using BIO RAD Clarity™ Western ECL substrate. BIO RAD ChemiDoc™ MP Imaging System and ImageLab 5.0 software were used to obtain the images.

### Annexin V/PI binding assay

Approximately 1 × 10^5^ number of cells were seeded in 24 well plate and treatments were done accordingly. Before starting the experiments, 1X filtered PBS were used to wash the cells twice. Next, according to the manufacturer’s protocol, cells were washed with 1X Annexin V/PI binding buffer and annexin V/FITC was added. PI was added at a concentration of 50 μg/ml and flow cytometry was performed by mixing the cells with 400 μl of 1X Annexin V binding buffer. The instrument for analysis, used was BIO-RAD S3e. Data was analyzed using FCS express (Licensed) and FLOWJO v10(Trail version).

### Gene Overexpression

Ectopic expressing of JNK was done using pcDNA3 Flag JNK2a1 (Addgene plasmid # 13754), transfection was performed with the help of Lipofectamine 3000 by following the manufacturer’s protocol. Post transfection, cells were treated with H_2_O_2_ 24 h and Western blot was performed to confirm the overexpression.

### Knock down assay

Approximately 2 × 10^6^ cells/well were plated in six-well plates and kept overnight in serum and antibiotics free optimal media. Next day, respective siRNA oligo was transfected into the cells by using Lipofectamine 3000 (Invitrogen, Carlsbad, California, USA). Based on dose–response studies the concentrations of siRNAs were chosen. After 48 h post transfection western blot was performed to check the knock down of the respective proteins.

### Co-immunoprecipitation

Approximately 3.5–4 million cells were used per treatment and washed with 1x PBS before proceeding for protein isolation. Next the cell pellet was dissolved in 100 μl of 1x RIPA and sonication was performed gently for 15–20 sec by keeping the sample on ice. The solution was then centrifuge at 15,000 rpm for 16 min at 4 ◦C and fresh 1.5 ml tube was used to collect the supernatant. After pre cleaning the cell lysate with IgG 200 µg of total protein was incubated overnight with primary antibody along with magnetic beads which was covalently linked using Pierce ^TM^ BS3 (LOT #XG341477) and next day the tubes were centrifuged at 1500 rpm for 10 min at 4 ◦C and the pellet were dissolved in 25 μl of sample buffer. Then the mixture was preheated at at 95 ◦C for 5 min and Western blot analysis was performed.

### Real time PCR

According to the manufacturer’s instructions, TRIZOL reagent, was used to isolate the RNA from the treated cells. Then agarose gel electrophoresis and Synergy H1 Microplate Reader was used to check the RNA quality and concentration respectively. iScript™ Reverse Transcription Supermix for RT-qPCR (BIO RAD) was used to prepare cDNA from isolated RNA. For Real-time PCR SsoFast™ Evagreen Supermix (BIO RAD), in CFX96 Touch™ Real-Time PCR Detection System (BIO RAD) was used. Minimum 40 cycles were run for each round of PCR for total saturation and the Cq values were considered for further analysis. All RT-qPCR setup were done in triplicate sets and the complete experiments were independently repeated at least three times. Actin was used as housekeeping gene for graphical representation of the gene expression data, and the mRNA levels were normalized accordingly.

### Immunofluorescences microscopy

Round glass bottom dish (Invitro Sci.) was used to seed the cells (0.2 to 0.4 million). Next cells were treated according to the standard protocol. After the treatment cells were fixed using 10 % PFA for 15-20 minutes. Washing was done using BSA containing NP-40 buffer for two hours before adding the primary antibodies (1:25) and incubated it for overnight. After incubation cells were again washed using NP-40 buffer for 2 hours and incubated with DAPI and secondary antibody (Alexa Fluro 594, CST) at 1:400 ratio prepared in NP-40 buffer. Before mounting the cells using mounting media cells were prewashed using the NP-40 wash buffer for two hours. All the images were obtained using Leica DMi8 fluorescence inverted microscope.

### Statistical Analysis

At least three independent experiments were performed before the data were analyzed as mean ±SD, and statistically significant differences amid mean values (from three independent experiment) were obtained using Student’s t-test.

## 4. Results

### 4.1 ROS mediated XIAP and FLIP downregulation takes different degradation pathways in drug sensitive and drug resistant leukemic cells

In our previous report we found that H_2_O_2_ downregulated both the IAP proteins FLIP and XIAP in a ubiquitin-proteasomal degradation pathway in K562R cells [13]. However, what is the fate of these proteins under ROS stress in K562S is still unknown. Therefore, both Imatinib resistant and sensitive K562 cells (K562R & K562S) were incubated overnight with indicated doses of H_2_O_2_ and from whole cell lysate western blot analysis was done using anti-FLIP and anti-XIAP antibodies. The results showed that FLIP and XIAP downregulated at the protein level dose dependently in both the sensitive (Fig. 1A) and resistant cells (Fig. 1B). Annexin V/PI assay with H_2_O_2_ indicated that H_2_O_2_ mediated apoptosis was not the reason for this reduction in XIAP and FLIP (Fig S1A). Real time qPCR showed there was no significant change at their transcript level in both K562S and K562R cells after H_2_O_2_ treatment (Fig. S1B). These results indicated that this downregulation of XIAP and FLIP by H_2_O_2_ might be via the protein degradation mechanisms. When we tried to identify the time points of these XIAP and FLIP downregulations, we found that XIAP downregulation start after 14 hours in K562S and after 12 hours in K562R cells but FLIP downregulation started after16 hours in K562S cells whereas it started after 14 hours in K562R cells (Fig. 1C, 1D). The onset of degradation at later time points in K562S cells indicated different degradation mechanisms for sensitive cells and resistant cell.

**Figure 1:**
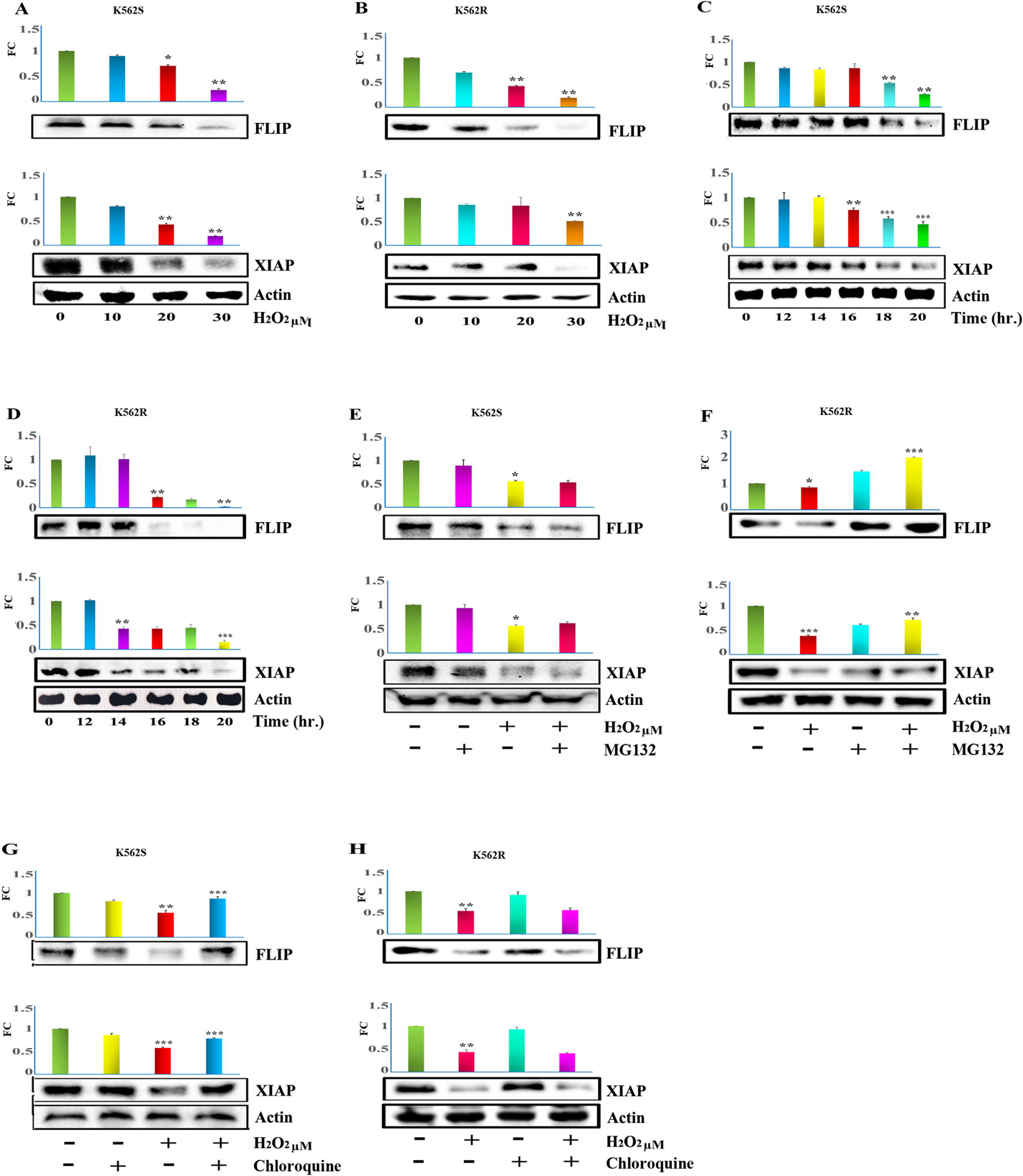
Downregulation of XIAP and FLIP via ROS takes different pathway in drug sensitive and drug resistant cells. A) Imatinib sensitive K562S cells were incubated with different doses of H_2_O_2_ overnight. Western blot analysis of FLIP and XIAP was made from whole cells lysate. As loading control γ Actin was used. Data were presented graphically as protein level fold change (FC) after densitometry analysis and indicate mean ± SD of three independent experiments. *p < 0.05, **p < 0.01, ***p < 0.001 B) Imatinib resistant K562R cells were incubated with different doses of H_2_O_2_ overnight. Western blot analysis of FLIP and XIAP was made from whole cells lysate. As loading control γ Actin was used. Data were presented graphically as protein level fold change (FC) after densitometry analysis and indicate mean ± SD of three independent experiments. *p < 0.05, **p < 0.01, ***p < 0.001 C) Time kinetics of FLIP and XIAP was measured using western blot analysis from imatinib sensitive K562 cells after treating the cells with 30 μM H_2_O_2_ for the specified time points. As loading control γ Actin was used. Data were presented graphically as protein level fold change (FC) after densitometry analysis and indicate mean ± SD of three independent experiments. *p < 0.05, **p < 0.01, ***p < 0.001 D) Time kinetics of FLIP and XIAP was measured using western blot analysis from imatinib resistant K562 cells after treating the cells with 30 μM H_2_O_2_ for the specified time points. As loading control γ Actin was used. Data were presented graphically as protein level fold change (FC) after densitometry analysis and indicate mean ± SD of three independent experiments. *p < 0.05, **p < 0.01, ***p < 0.001 E) K562S cells were treated with either 30 μM H_2_O_2_ alone or in combination of 1.5 μM of MG132 for 16 h and the whole cell lysates were subjected to western blot analysis for XIAP and FLIP. Data were presented graphically as relative fold change (RF) after densitometry analysis and indicate mean ± SD of three independent experiments. *p < 0.05, **p < 0.01, ***p < 0.001 F) K562R cells were treated with either 30 μM H_2_O_2_ alone or in combination of 1.5 μM of MG132 for 16 h and the whole cell lysates were subjected to western blot analysis for XIAP and FLIP. As loading control γ Actin was used. Data were presented graphically as relative fold change (RF) after densitometry analysis and indicate mean ± SD of three independent experiments. *p < 0.05, **p < 0.01, ***p < 0.001 G) K562S cells were treated with 30 μM H_2_O_2_ either alone or in combination with 20 μM Chloroquine for 16 h and the whole cell lysates were subjected to Immunoblot analysis with anti-FLIP and anti-XIAP antibodies. As loading control γ Actin was used. Data were presented graphically as protein level fold change (FC)after densitometry analysis and indicate mean ± SD of three independent experiments. *p < 0.05, **p < 0.01, ***p < 0.001 H) K562R cells were treated with 30 μM H_2_O_2_ either alone or in combination with 20 μM Chloroquine for 16 h and the whole cell lysates were subjected to Immunoblot analysis with anti-FLIP and anti-XIAP antibodies. As loading control γ Actin was used. Data were presented graphically as protein level fold change (FC) after densitometry analysis and indicate mean ± SD of three independent experiments. *p < 0.05, **p < 0.01, ***p < 0.001

One of the major post translational modification of proteins is ubiquitination and degradation via lysosomal or proteasomal pathways [14],[15]. Therefore, Involvement of these two pathways were analyzed. Proteasome inhibitor MG132 [16] and lysosome pathway inhibitor chloroquine [17] were separately pretreated in K562S and K562R cells prior to the treatment with H_2_O_2_. Western blot analysis showed that, MG132, the proteasome inhibitor, could not reverse the H_2_O_2_ mediated degradation of XIAP and FLIP in drug sensitive CML cells (Fig. 1E), whereas, it reversed the same in drug resistant CML cells (Fig. 1F). In contrast, autophagy inhibitor Chloroquine reversed the degradation of XIAP and FLIP by H_2_O_2_ in drug sensitive cells (Fig. 1G) but failed to do so in drug resistant cells (Fig. 1H).

### 4.2 XIAP and FLIP degradation takes ubiquitin mediated autophagic pathway in drug sensitive cells

The involvement of lysosomal pathway was further checked by immunofluorescence after K562(S) cells were treated with both MG132 and Chloroquine prior to treatment of H_2_O_2_ (Fig. 2A, B). The results confirmed that lysosomal inhibitor Chloroquine but not proteasomal inhibitor MG132 reversed both FLIP and XIAP degradation in K562(S) cells. Ubiquitin ligase plays an important role in all the ubiquitination process and these ubiquitin ligases are very specific for their targets. Literature suggests that ITCH is an E3 ubiquitin ligase enzyme which plays a critical role in cellular protein ubiquitination and degradation via proteasomal and lysosomal pathways [18,19]. Immunoblot analysis revealed increased ubiquitin smear band (Fig. 2C, D) when K562(S) cells were treated with Chloroquine prior to the treatment with H_2_O_2_, then western blot with anti-ubiquitin antibody was performed with anti-ubiquitin antibody after immunoprecipitation with anti-FLIP and anti-XIAP antibody, separately. These clearly indicated ubiquitination of XIAP and FLIP under H_2_O_2_ treatment. Since all these results confirmed the autophagy-mediated degradation in K562S cells and proteasome mediated degradation for K562R cells, we checked the global autophagy status of both drug sensitive and drug resistant cells under the H_2_O_2_ treatment. p62 and LC3BII, two well-known autophagy markers were checked in K562S and K562R cells with H_2_O_2_. Interestingly the outcomes confirmed that, under the ROS treatment in drug sensitive cell, the p62 level was decreased and LC3BII level increased (Fig. 2E) and in drug resistant cells both p62 and LC3BII remained unaltered (Fig. 2E). All of these data confirmed that the degradation of XIAP and FLIP were via proteasomal pathway in K562R cells and via lysosomal pathway in K562S cells. Moreover, it was also confirmed that global autophagy level remained upregulated under the ROS treatment in drug sensitive cells but not in drug-resistant cells. Now, ITCH mediated ubiquitination and degradation of FLIP and autoubiquitination and degradation of XIAP was previously reported by us in our previous study in K562R cells [20]. Here also when immunoprecipitation was performed, under H2O2 treatment, using anti-ITCH antibody and then immunoblotted using anti XIAP and FLIP antibody, the results showed that ITCH indeed binds with FLIP and ITCH level increased upon H_2_O_2_ treatment (Fig. 2F left panel). However, XIAP neither bound to ITCH nor this binding has any role of H_2_O_2_ (Fig. 2F right panel).

**Figure 2:**
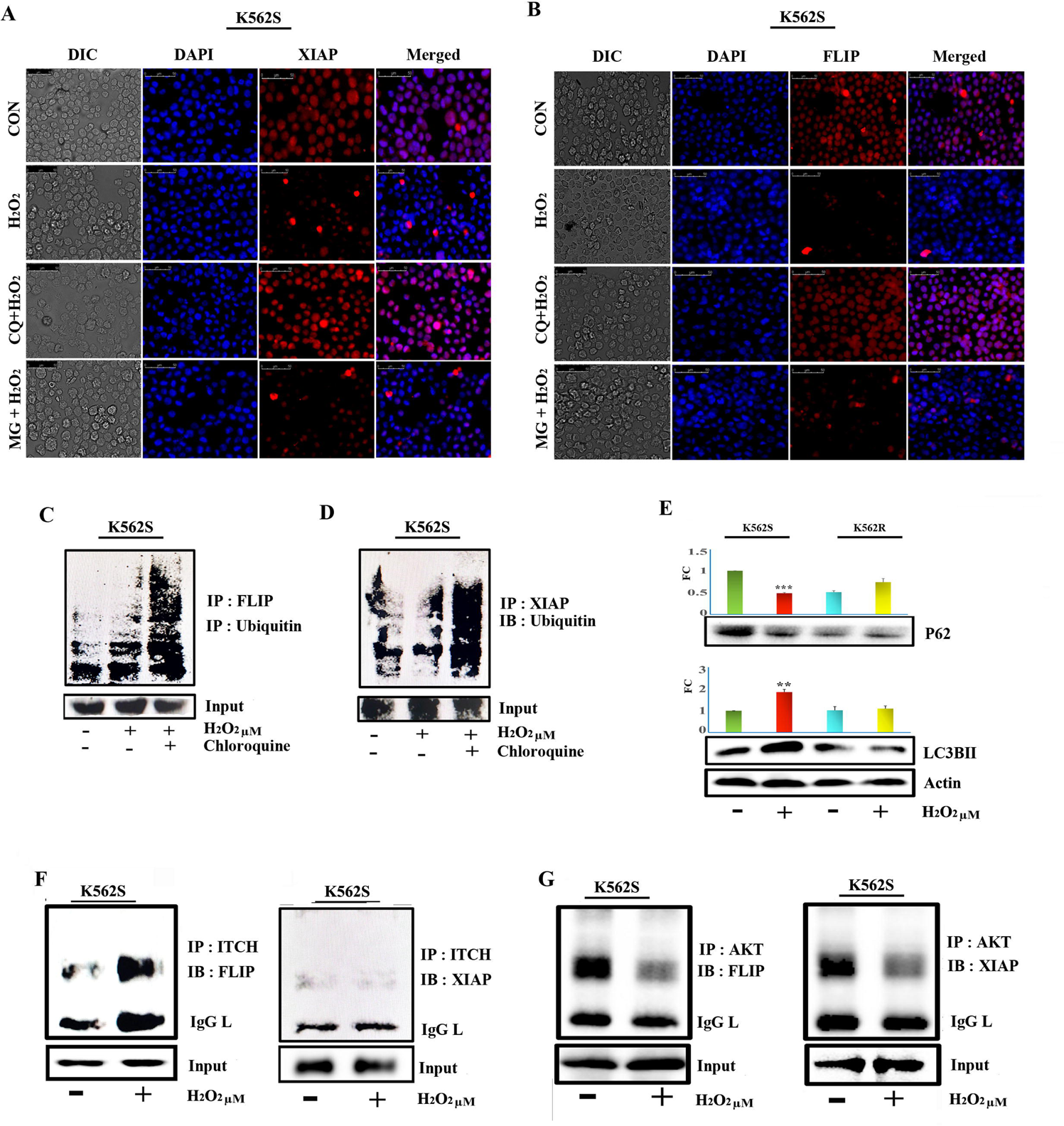
XIAP and FLIP degradation takes ubiquitin mediated autophagic pathway in drug sensitive cells. A) K562S cells were treated with 30 μM H_2_O_2_ alone or in combination 1.5 μM of MG132 and in combination with 20 μM Chloroquine for 14 hours, and the cells were fixed using 10% PFA onto the slides and incubated with anti XIAP antibody overnight and stained with DAPI (Blue) and Alexa Fluro 594 (Red). B) K562S cells were treated with 30 μM H_2_O_2_ alone or in combination 1.5 μM of MG132 and in combination with 20 μM Chloroquine for 14 hours, and the cells were fixed using 10% PFA onto the slides and incubated with anti-FLIP antibody overnight and stained with DAPI (Blue) and Alexa Fluro 594 (Red). C) K562S cells were treated with 30 μM H_2_O_2_ alone or in combination with 20 μM Chloroquine for 14 h and the whole cell lysates were subjected to co-Immunoprecipitation with FLIP antibody and then Western blot analysis were performed with anti-ubiquitin antibody. The lower panel shows the input of FLIP and upper panel, the immunoprecipitates. D) K562S cells were treated with 30 μM H_2_O_2_ alone or in combination with 20 μM Chloroquine for 14 h and the whole cell lysates were subjected to co-Immunoprecipitation with XIAP antibody and then Western blot analysis were performed with anti-ubiquitin antibody. The subordinate panel shows the input of XIAP and upper panel, the immunoprecipitates. E) K562S and K562R cells were treated with 30 μM H_2_O_2_ alone for 12-13 hours and the whole cell lysates were subjected to western blot analysis using anti P62 and LC3BII antibody. As loading control γ Actin was used. Data were presented graphically as protein level fold change (FC) after densitometry analysis and indicate mean ± SD of three independent experiments. *p < 0.05, **p < 0.01, ***p < 0.001 F) K562(S) cells were treated with or without H_2_O_2_ for 14 hours and subjected to immunoprecipitation using anti-ITCH antibody and then immunoblotted using anti-FLIP and anti-XIAP antibody. FLIP and XIAP was used as input. The data is representative of three independent experiments. G) K562(S) cells were treated with 30 μM H2O2 for 16 hours and whole cell lysate was subjected to co-immunoprecipitation using anti AKT antibody and then immunoblotted using anti-FLIP and anti-XIAP antibody. FLIP and XIAP was used as input. The data is representative of three independent experiments.

Now, according to the previous findings in ovarian cancer cells, AKT upon phosphorylation provide a structural and functional stability to both XIAP and FLIP [21,22]. If this is true for K562S cells, then there will be a change in physical interaction between XIAP and Akt prior to its degradation. We previously showed that XIAP binds to Akt under the influence of H_2_O_2_ [13]. Therefore, K562(S) cells were treated for 16 hours with 30 μM H_2_O_2_ and whole cell lysate was used to perform co-immunoprecipitation using anti AKT antibody and then immunoblotted using anti XIAP antibody. Results showed that Akt indeed bound to both XIAP and ROS treatment reduced the binding of AKT with XIAP (Fig. 2G). Very interestingly, when we did the same experiment and immunoblotted FLIP after immunoprecipitation with Akt antibody, it also revealed physical interaction of Akt and FLIP and this interaction decreased upon H_2_O_2,_ treatment. Therefore, it was worth searching, at this stage, whether PI3k/Akt pathway has any role in both of these XIAP and FLIP downregulation.

### 4.3 Involvement of AKT-mTOR pathway in H_2_O_2_ mediated FLIP and XIAP degradation

It was confirmed from our data that, XIAP and FLIP down regulation mediated via lysosomal pathway under H_2_O_2_ treatment in drug sensitive cells, whereas UPS (ubiquitin proteasome system) control the overall degradation under H_2_O_2_ treatment in drug resistant cells. Furthermore, we have found that, AKT dissociate from both the XIAP and FLIP under the ROS treatment in K562R cells. Thus, we checked the activation of PI3K/Akt pathway. It is a well-known fact that phosphorylation of AKT is needed for its activation and PI3K, upon phosphorylation and activation, in turn, activates AKT. Western blot analysis showed H_2_O_2_ dose dependently reduces both PI3K and Akt phosphorylation, whereas the basal level of both AKT and PI3K remained unaltered under the ROS treatment (Fig. 3A). Now, to identify the role of AKT/PI3K in the downregulation of XIAP and FLIP, K562(S) cells were incubated with different doses of PI3K inhibitor wortmannin for 14 hours and then western blot analysis with the whole cell lysates showed that wortmannin dose dependently downregulated both XIAP and FLIP in K562S cells (Fig. 3B) and Wortmannin in combination with H_2_O_2_ decreased the FLIP and XIAP more than H_2_O_2_ alone (Fig 3C).

**Figure 3:**
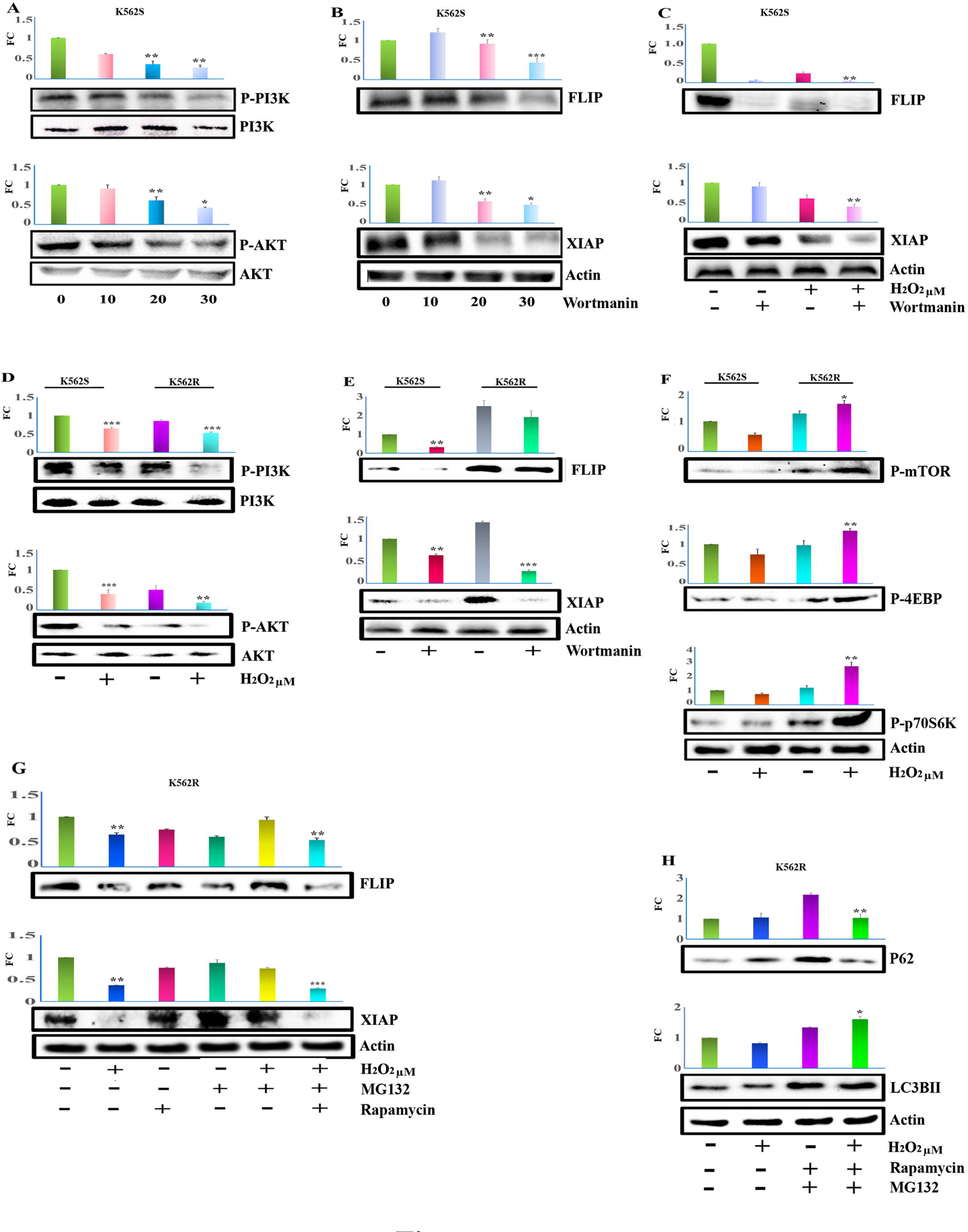
Involvement of AKT-mTOR pathway in H_2_O_2_ mediated FLIP and XIAP degradation. A) K562(S) cells were treated with indicated dose of H_2_O_2_ and whole cell lysate were subjected to western blot analysis using AKT, pAKT and PI3K, pPI3K antibodies. γ Actin was used as loading control. Data were presented graphically as protein level fold change (FC) after densitometry analysis and indicate mean ± SD of three independent experiments. *p < 0.05, **p < 0.01, ***p < 0.001 B) K562(S) cells were treated with different doses of PI3K inhibitor wortmannin for 14 hours, whole cell lysate was subjected to western blot analysis using anti-FLIP and anti-XIAP antibodies. γ Actin was used as loading control. Data were presented graphically as protein level fold change (FC) after densitometry analysis and indicate mean ± SD of three independent experiments. *p < 0.05, **p < 0.01, ***p < 0.001 C) K562(S) cells were pre-treated with PI3K inhibitor Wortmannin prior to the treatment of 30 μM H2O2 for 16 hours, and whole cell lysate were subjected to western blot using anti XIAP and FLIP antibody. As loading control γ Actin was used. Data were presented graphically as protein level fold change (FC) after densitometry analysis and indicate mean ± SD of three independent experiments. *p < 0.05, **p < 0.01, ***p < 0.001 D) Both K562(S) and K562(R) cells were treated 30 μM H2O2 for 16 hours and whole cell lysate were subjected to western blot analysis using AKT, pAKT and PI3K, pPI3K antibodies. γ Actin was used as loading control. Data were presented graphically as protein level fold change (FC) after densitometry analysis and indicate mean ± SD of three independent experiments. *p < 0.05, **p < 0.01, ***p < 0.001 E) Both K562(S) and K562(R) cells were treated with PI3K inhibitor wortmannin (30 μM) for 16 hours and whole cell lysate were subjected to western blot analysis using anti FLIP and XIAP antibodies. γ Actin was used as loading control. Data were presented graphically as protein level fold change (FC) after densitometry analysis and indicate mean ± SD of three independent experiments. *p < 0.05, **p < 0.01, ***p < 0.001 F) Both K562(S) and K562(R) cells were treated with 30 μM H_2_O_2_ and whole cell lysate were subjected to western blot analysis using phospho mTOR, phospho 4EBP and phospho P70S6K, antibodies. γ Actin was used as loading control. Data were presented graphically as protein level fold change (FC) after densitometry analysis and indicate mean ± SD of three independent experiments. *p < 0.05, **p < 0.01, ***p < 0.001 G) K562(R) cells were treated with 1.5 μM of MG132 and 200 nM of rapamycin prior to the treatment with 30 μM H_2_O_2_ for 16 hours and whole cell lysate were subjected to western blot analysis using XIAP and FLIP antibodies. γ Actin was used as loading control. Data were presented graphically as protein level fold change (FC) after densitometry analysis and indicate mean ± SD of three independent experiments. *p < 0.05, **p < 0.01, ***p < 0.001 H) K562(R) cells were treated with 200 nM of Rapamycin prior to the treatment with 30 μM H_2_O_2_ and whole cell lysate were subjected to western blot analysis using P62 and LC3BII antibodies. γ Actin was used as loading control. Data were presented graphically as protein level fold change (FC) after densitometry analysis and indicate mean ± SD of three independent experiments. *p < 0.05, **p < 0.01, ***p < 0.001

Moreover, Western blot analysis with the whole cell lysates from H_2_O_2_-treated K562S and K562R revealed that H_2_O_2_ decreased the phosphorylation of PI3K and Akt in both sensitive and resistant cells (Fig 3D). Interestingly, the inherent phospho-PI3K and Phospho-Akt were observed to be way higher in sensitive cells than in resistant cells (Fig 3D). Similarly, comparison of K562S and K562R cells with wortmannin treatment revealed that wortmannin could decrease XIAP level in both the sensitive and resistant cells but it could decrease FLIP only in K562S cells (Fig 3E). These data clearly revealed that PI3K/Akt has different influence on XIAP and FLIP downregulation in K562R and K562S cells and thus it could be assumed that the signaling pathways that lead to the activation of these degradation pathways might be different for K562S and K562R.

Now, since both PI3K and Akt phosphorylation decreased in the same way in K562S and K562R cells but XIAP and FLIP degradation patterns are different in these two cells, the downstream targets were considered worth to check. The mTOR is a very well-known downstream target of PI3K/Akt, especially in case of any kind of stress. Thus, we checked the activation of mTOR pathway in drug resistant and sensitive cells. H_2_O_2_ treatment for 12 hours and subsequent western blot analysis showed that phospho-mTOR, phospho-4EBP and phospho-P70S6K levels were increased upon H_2_O_2_ treatment in drug resistant cells only (Fig. 3F). Not only that, the endogenous phosphorylation level of all these proteins were observed higher in resistant cells than in the sensitive cells. Now, if, mTOR activation was responsible for proteasomal degradation of FLIP and XIAP in K562R cells, then inhibition of mTOR should result in XIAP and FLIP degradation even after proteasomal pathway inhibition. Therefore, Drug resistant cells were treated with both rapamycin and MG132 before H_2_O_2_ treatment for 16 hours. Western blot analysis revealed that the H_2_O_2_ treatment is involved in the degradation of XIAP and FLIP when both mTOR and proteasomal pathway are inhibited by Rapamycin and MG132 respectively (Fig. 3G). However the degradation reversed when MG132 alone was pre-treated before H_2_O_2_ addition. To confirm the role of mTOR, we treated the drug resistant cells with both mTOR inhibitor rapamycin and Proteasome inhibitor MG132 and then incubated with H_2_O_2_ for 12 hours. Western blot analysis using anti p62, anti LC3BII antibodies revealed that P62 level was reduced and the amount of LC3BII were increased in only where both the inhibitors were pre-treated (Fig. 3H) which confirmed that H_2_O_2_ and rapamycin treatment activated the autophagy pathway in the K562R cells. Thus, it is clear from our data that in K562(R) cells PI3K is involved in XIAP downregulation but not FLIP and in K562(S) cells PI3K is involved in both XIAP and FLIP downregulation and mTOR played a crucial role in these differential regulations. Thus, all these data conclusively told that XIAP and FLIP downregulation via proteasomal pathways is the result of mTOR pathway activation in K562R cells. The degradation would have taken lysosomal pathway had the mTOR pathway been remained inhibited. Therefore, we tried to find out why mTOR activation resulted in and XIAP and FLIP degradation in K562R cells and what are the difference in the upstream signaling between K562R and K562S cells.

### 4.4 Differential activation of MAPK pathway in drug sensitive and drug resistant cells by H_2_O_2_

Previously we showed that ROS mediated downregulation of XIAP and FLIP was depended on MAP kinase pathway in K562R cells[13]. We also showed that ROS activated ERK downregulated XIAP and activation JNK by ROS was necessary for the downregulation of FLIP in K562R cells[13]. Thus, here we have checked whether this MAP kinase pathway components like JNK, p38, ERK are involved in the mTOR activation and thus activation of differential degradation pathway. Western blot analysis indicated that both XIAP and FLIP downregulation by ROS was significantly reversed by JNK inhibitor SP (21) (Fig. 4A). However, ERK inhibitor PD (22,23) did not reverse any of the XIAP and FLIP downregulation by H_2_O_2_ (Fig. 4B). We have repeated the experiment with JNK and ERK inhibitors in K562R cells and western blot analysis with cell lysates showed that FLIP downregulation by H_2_O_2_ was reversed by JNK inhibitor but not by ERK inhibitor whereas XIAP downregulation was significantly reversed by ERK inhibitor but not by JNK inhibitor in K562R cells (Fig. 4C). This confirmed that downregulation of both XIAP and FLIP is dependent on JNK activation only in drug sensitive cells, whereas, in K562R cells, JNK activations were involved in FLIP and ERK was involved in XIAP downregulations. When we checked the phosphorylation status of JNK with H_2_O_2_ in K562S cells, western blot analysis showed that H_2_O_2_ dose dependently amplified JNK phosphorylation keeping its protein level unaltered (Fig. 4D). However, interestingly, ERK phosphorylation was found to be significantly decreased dose dependently upon H_2_O_2_ treatment without any significant change in protein level in K562S cells (Fig. 4E) whereas, H_2_O_2_ treatment dose dependently increased both JNK and ERK phosphorylation in K562R cells (Fig. 4F). When similar experiments were performed to check the other component of MAP kinase pathway p38, the phosphorylation of p38 and its protein level did not show any change (Supplementary Figure S1, C). Moreover, Phosphorylation of JNK was checked at different time points by treating the K562(S) cells with 30μM H2O2. The results indicated that JNK phosphorylation increased from 4 hours and reached its peaked at 16-hour time point (Fig. S1D) which perfectly correlated with the FLIP and XIAP downregulation time which also indicated that both XIAP and FLIP degradation is mediated by activation of JNK alone. The most interesting observation here in these results is that ERK was inhibited by H_2_O_2_ in the K562S cells whereas it got activated by H2O2 in K562R cells. Is this activation played a role in mTOR activation and subsequent lysosomal pathway inhibition in K562R cells?

**Figure 4:**
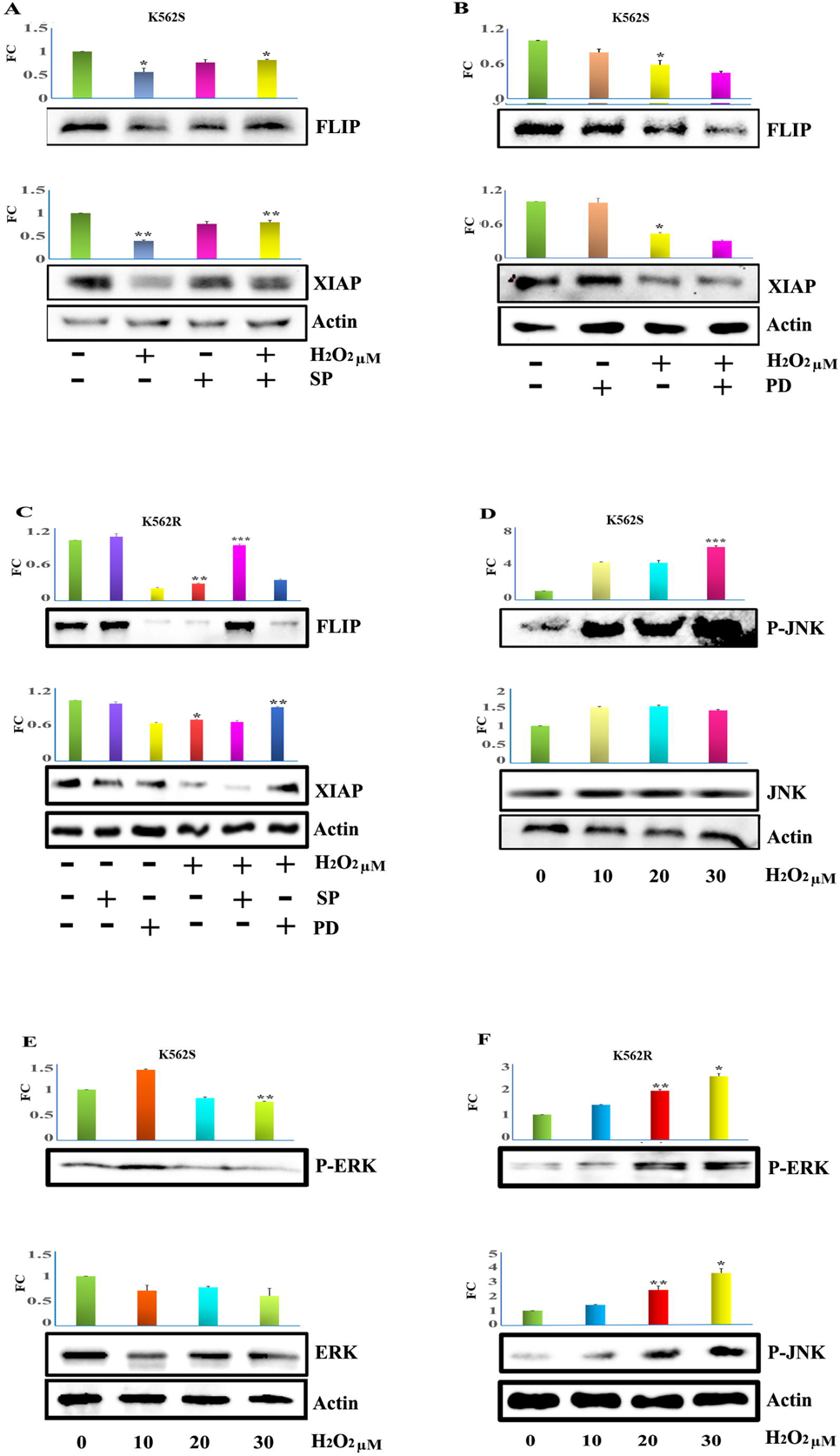
Differential activation of MAPK pathway in drug sensitive and drug resistant cells by H_2_O_2_. A) K562(S) cells were treated with JNK inhibitor SP600125(21) prior to the incubation with H_2_O_2_ and cell lysates were subjected to western blot analysis with anti XIAP and FLIP antibody. γ Actin was used as loading control. Data were presented graphically as protein level fold change (FC) after densitometry analysis and indicate mean ± SD of three independent experiments. *p < 0.05, **p < 0.01, ***p < 0.001 B) K562(S) cells were treated with ERK inhibitor PD98059(22,23) prior to the incubation with H_2_O_2_ and cell lysates were subjected to western blot analysis with anti XIAP and FLIP antibody. γ Actin was used as loading control. Data were presented graphically as protein level fold change (FC) after densitometry analysis and indicate mean ± SD of three independent experiments. *p < 0.05, **p < 0.01, ***p < 0.001 C) K562R cells were treated with JNK inhibitor SP600125(21) and ERK inhibitor PD98059(22,23) prior to the incubation with H_2_O_2_ and cell lysates were subjected to western blot analysis with anti XIAP and FLIP antibody. As loading control γ Actin was used. Data were presented graphically as protein level fold change (FC) after densitometry analysis and indicate mean ± SD of three independent experiments. *p < 0.05, **p < 0.01, ***p < 0.001 D) K562(S) cells were incubated with different doses of H_2_O_2_ for 14 hour and whole cell lysate were subjected to western blot analysis with anti JNK and anti p-JNK antibodies. γ Actin was used as loading control. Data were presented graphically as protein level fold change (FC) after densitometry analysis and indicate mean ± SD of three independent experiments. *p < 0.05, **p < 0.01, ***p < 0.001 E) K562(S) cells were incubated with different doses of H_2_O_2_ for 14 hour and whole cell lysate were subjected to western blot analysis with anti ERK and anti p-ERK antibodies. γ Actin was used as loading control. Data were presented graphically as protein level fold change (FC) after densitometry analysis and indicate mean ± SD of three independent experiments. *p < 0.05, **p < 0.01, ***p < 0.001 F) K562(R) cells were incubated with different doses of H_2_O_2_ for 14 hour and whole cell lysate were subjected to western blot analysis with anti p-ERK and p-JNK antibodies. γ Actin was used as loading control. Data were presented graphically as protein level fold change (FC) after densitometry analysis and indicate mean ± SD of three independent experiments. *p < 0.05, **p < 0.01, ***p < 0.001

### 4.5 Genetic manipulation of JNK and ERK influences the XIAP and FLIP degradation

We have previously found that both phospho-ERK level and phospho-mTOR pathway were upregulated under the H_2_O_2_ treatment in drug resistant cells. Thus, we tried to identify whether ERK pathway has any influence on mTOR pathway. ERK was knocked down in drug resistant cells using ERK Si-RNA and cells were treated with H_2_O_2_ for 12 hours. Western blot analysis showed knocked down of ERK reversed the increased phospho-mTOR level significantly even after H_2_O_2_ treatment (Fig. 5A). Next, we checked how these ERK knockdown influence the autophagy activation. when we knocked down ERK using ERK siRNA and checked the autophagy status upon H_2_O_2_ treatment, we found that H_2_O_2_ treatment decreased p62 level and incresed LC3BII level, indicating activation of lysosomal degradation pathway in K562R cells (Fig. 5B). Thus, these data confirmed that ERK mediated mTOR activation played a critical role in the suppression of autophagy pathway in K562R cells. Since ERK phosphorylation decreased with the treatment of H_2_O_2_ in K562S cells, the phospho-mTOR remains inhibited and autophagy remined activated in K562S cells.

**Figure 5:**
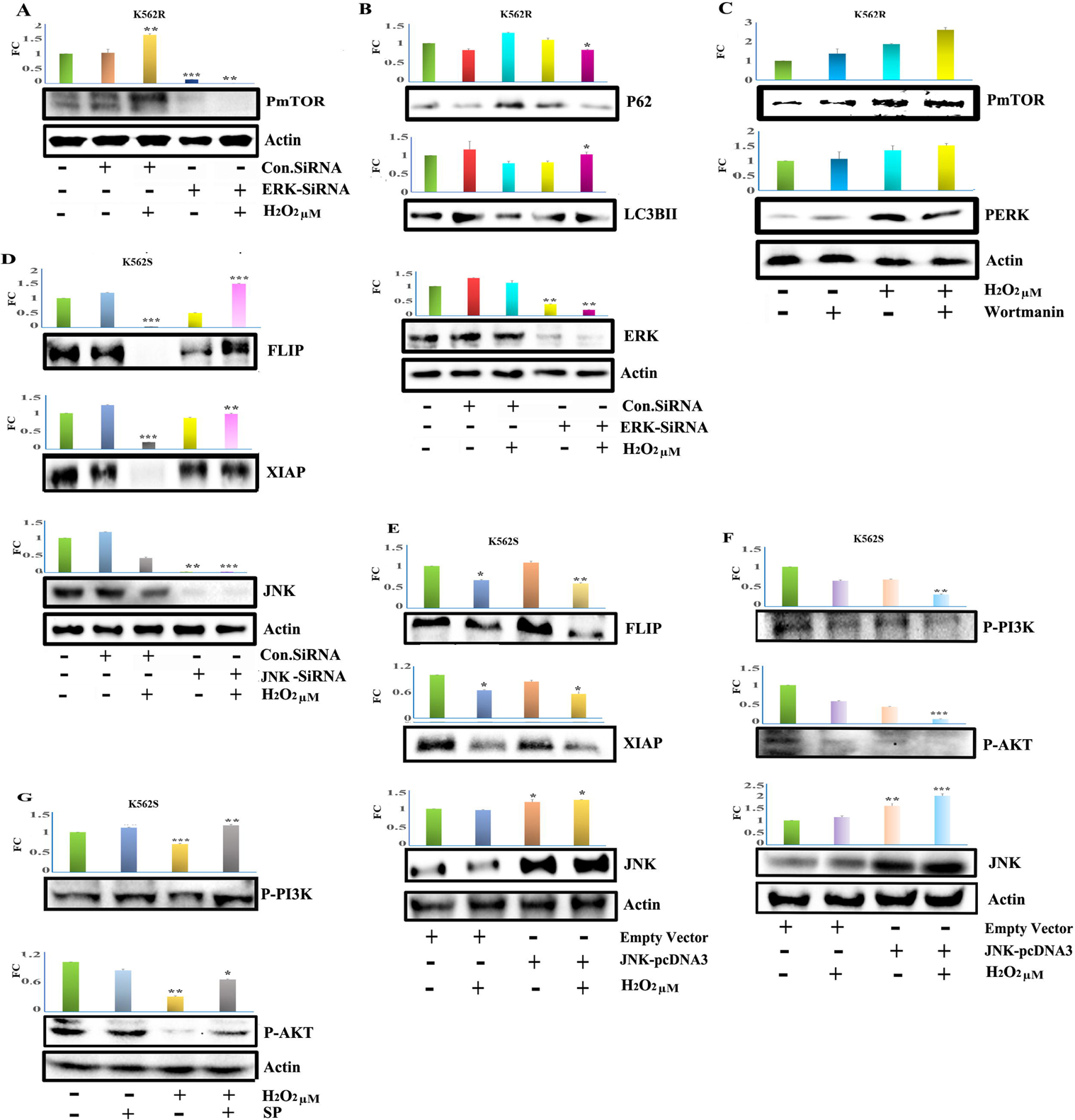
Genetic manipulation of JNK and ERK influences the XIAP and FLIP degradation. A) Knocked down of ERK was achieved by using siRNA in K562(R) cells and immunoblot analysis of phospho mTOR were performed with the whole cell lysates taken after 14 h of 30 μM H_2_O_2_ treatment. Scrambled siRNA was used as control. Data were presented graphically as protein level fold change (FC) after densitometry analysis and indicate mean ± SD of three independent experiments. *p < 0.05, **p < 0.01, ***p < 0.001 B) Knocked down of ERK was achieved by using siRNA in K562(R) cells and immunoblot analysis of P62, LC3BII and ERK were performed with the whole cell lysates taken after 14 h of 30 μM H_2_O_2_ treatment. Scrambled siRNA was used as control. Data were presented graphically as protein level fold change (FC) after densitometry analysis and indicate mean ± SD of three independent experiments. *p < 0.05, **p < 0.01, ***p < 0.001 C) K562(R) cells were pre-treated with PI3K inhibitor Wortmannin prior to the treatment of 30 μM H2O2 for 16 hours, and whole cell lysate were subjected to western blot using anti phospho-mTOR and phospho-ERK antibody. As loading control γ Actin was used. Data were presented graphically as protein level fold change (FC) after densitometry analysis and indicate mean ± SD of three independent experiments. *p < 0.05, **p < 0.01, ***p < 0.001 D) Knocked down of JNK was achieved by using siRNA in K562(S) cells and immunoblot analysis of FLIP, XIAP and JNK were performed with the whole cell lysates taken after 16 h of 30 μM H_2_O_2_ treatment. As loading control γ Actin was used. Data were presented graphically as protein level fold change (FC) after densitometry analysis and indicate mean ± SD of three independent experiments. *p < 0.05, **p < 0.01, ***p < 0.001 E) K562(S) cells were transfected with either pcDNA3 Flag JNK2a1 or empty vector and Immunoblot analysis of FLIP, XIAP and JNK were performed after 30 μM of H_2_O_2_ treatment for 14 h. As loading control γ Actin was used. Data were presented graphically as protein level fold change (FC) after densitometry analysis and indicate mean ± SD of three independent experiments. *p < 0.05, **p < 0.01, ***p < 0.001 F) pcDNA3 Flag JNK2a1 or empty vector were transfected in K562(S) cells and Immunoblot analysis was performed of pPI3K and pAKT and JNK after 30 μM of H_2_O_2_ treatment for 14 h. As loading control γ Actin was used. Data were presented graphically as protein level fold change (FC) after densitometry analysis and indicate mean ± SD of three independent experiments. *p < 0.05, **p < 0.01, ***p < 0.001 G) K562S cells were treated with JNK inhibitor SP600125(21) prior to the incubation with H_2_O_2_ and cell lysates were used for western blot analysis with anti pPI3K and pAKT antibodies. As loading control γ Actin was used. Data were presented graphically as protein level fold change (FC) after densitometry analysis and indicate mean ± SD of three independent experiments. *p < 0.05, **p < 0.01, ***p < 0.001

Since PI3K/Akt pathways are known to be involved in mTOR signaling, we tested the involvement of this pathway. Therefore, K562(R) cells were pre-treated with PI3K inhibitor Wortmannin prior to the treatment of H_2_O_2_ and whole cell lysate were used for western blot analysis using anti phospho-mTOR and phospho-ERK antibody. Results indicated PI3K does not play any role in this phospho-ERK mediated increased m-TOR phosphorylation (Fig. 5C).

In figure 4, we have found that JNK activation played a critical role in both XIAP and FLIP downregulation in K562S cells. To further confirm whether only JNK activation is sufficient for the H_2_O_2_ mediated downregulation of XIAP and FLIP, JNK was knocked down using JNK siRNA in K562(S) cells and treated with 30 μM H_2_O_2_ for 16 hours and whole cell lysate was used for western blot analysis using anti XIAP and FLIP antibodies. Scrambled siRNA was used as control. Results indicated that knock down of JNK reversed the downregulation of XIAP and FLIP significantly (Fig. 5D). To confirm our hypothesis, JNK was ectopically over expressed by transfecting the K562(S) cells with pcDNA3 Flag JNK2a1 plasmid and then cells were treated with and without 30 μM H_2_O_2_ for 16 hours. Western blot analysis was performed using whole cell lysate with anti XIAP, FLIP showed that pcDNA3 Flag JNK2a1 plasmid containing cell showed more XIAP & FLIP downregulation compared to empty vector transfected cells when treated with H_2_O_2_ (Fig. 5E). Now, we have already shown that PI3K and AKT has an important role to play[23,24], therefore, it was checked whether PI3K and Akt are controlled by JNK to exert its effect on XIAP and FLIP. JNK over expressed K562(S) cells were treated with or without 30 μM H_2_O_2_ for 14 hours and whole cell lysate were used for western blot analysis using and anti-phospho PI3K and phospho AKT antibodies. Results displayed that upon over expression of JNK, both phospho PI3K and phospho AKT level were reduced significantly when K562(S) cells were treated with H_2_O_2_ (Fig. 5F). On the other hand, we also showed that in presence of JNK inhibitor SP600125 H_2_O_2_ mediated dephosphorylation of PI3K and AKT were reversed (Fig. 5G). Thus, all these data confirmed that JNK activation alone was sufficient to decrease XIAP and FLIP via PI3K/Akt inactivation in K562S cells. Moreover, it was also confirmed that mTOR activation was the result of Erk activation in K562R cells and thus Erk played a significant role in XIAP and FLIP degradation via proteasomal pathway by suppressing the autophagy.

### 4.6 Phospho-mTOR is responsible for Imatinib resistance and enhanced ROS level in K562R cells

Now, we have found that H_2_O_2_ differentially regulate XIAP and FLIP degradation in two different degradation pathways in K562(R) and K562(S) cells by alteration in the upstream signaling molecules like PI3K/Akt, mTOR and ERK. Therefore, we checked the inherent endogenous ROS level in K562(R) and K562(S) cells. K562S and K562R cells were stained with DCFDA and analyzed in flow cytometer. We found that K562R cells contains almost double endogenous ROS than the K562S cells (Fig. 6A). It is worth to mention here that we have already found that mTOR remained activated in K562R cells. Therefore, we tried to check if there is any link between these enhanced ROS level and mTOR activation. Interestingly, western blot analysis revealed that ROS scavenger NAC, indeed, reversed that higher phosphorylation level of mTOR (Fig 6B). Therefore, it may be concluded that higher endogenous mTOR activation might be due to higher ROS level in K562R cells.

**Figure 6:**
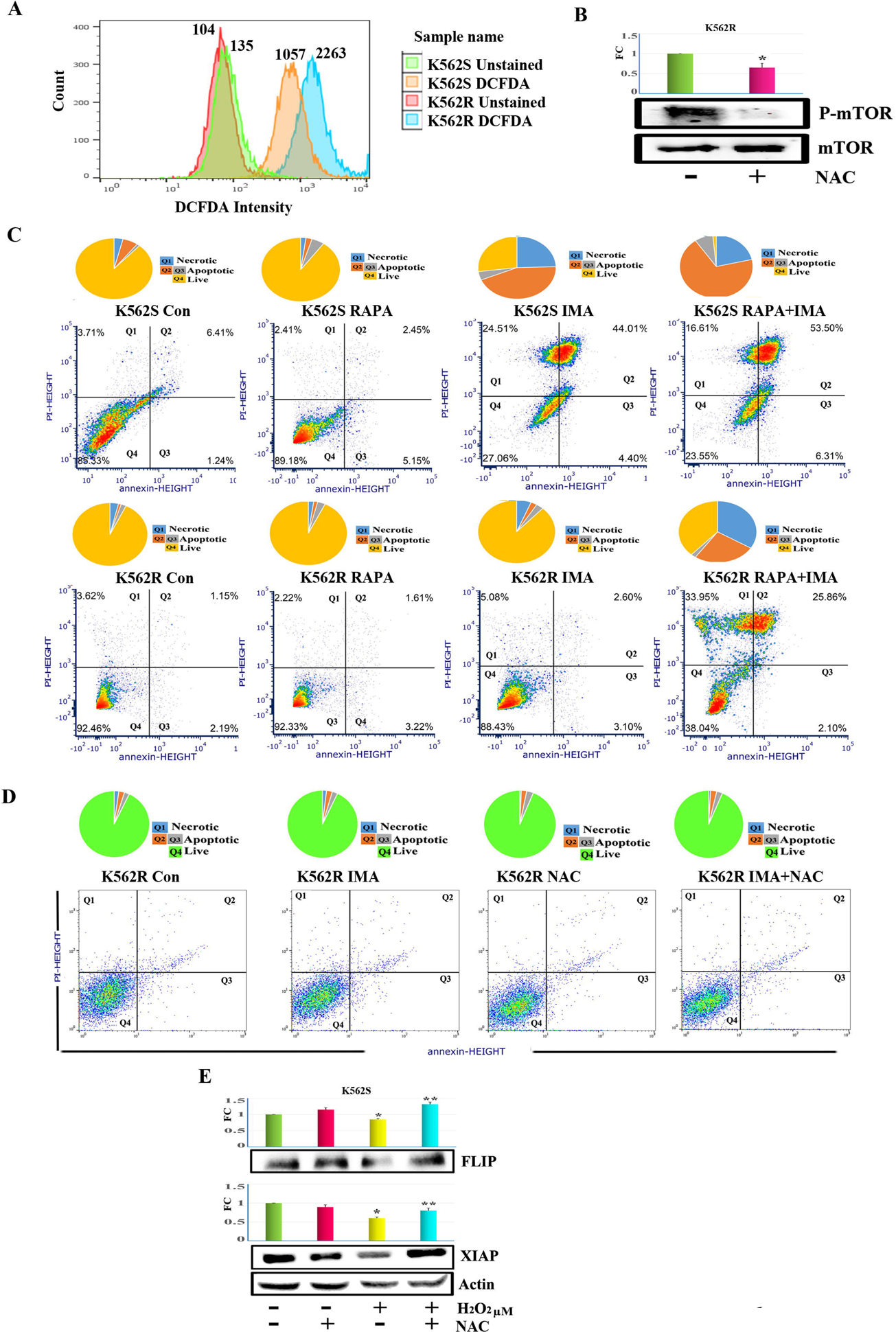
Phospho-mTOR is responsible for Imatinib resistance and enhanced ROS level in K562R cells. A) Both K562(S) and K562(R) cells were stained with DCFDA and subjected for flowcytometry analysis to measure the indigenous ROS. The data is representative of three independent experiments. B) Both K562(S) and K562(R) cells were treated with rapamycin alone or with imatinib (2 μM) overnight and subjected for flowcytometry analysis of Annexin V/PI assay. The data is representative of three independent experiments. The data is representative of three independent experiments. Top panel is the graphical illustration of several population in pie chart and lower panel denotes demonstrative dot plots. C) K562(R) cells were treated with NAC (1mM) alone or with imatinib (2 μM) overnight and subjected for flowcytometry analysis of Annexin V/PI assay. The data is representative of three independent experiments. Top panel is the graphical illustration of several population in pie chart and lower panel denotes demonstrative dot plots. D) K562S cells were treated with ROS scavenger NAC alone and in combination with H_2_O_2_, cell lysate was subjected to western blot analysis using anti FLIP and XIAP antibodies. Data were presented graphically as protein level fold change (FC) after densitometry analysis and indicate mean ± SD of three independent experiments. *p < 0.05, **p < 0.01, ***p < 0.001 E) K562(R) cells were treated with NAC (1mM) alone overnight, cell lysates were used for western blot analysis with anti mTOR and phospho-mTOR antibodies. Data were presented graphically as protein level fold change (FC) after densitometry analysis and indicate mean ± SD of three independent experiments. *p < 0.05, **p < 0.01, ***p < 0.001

**Figure 7:**
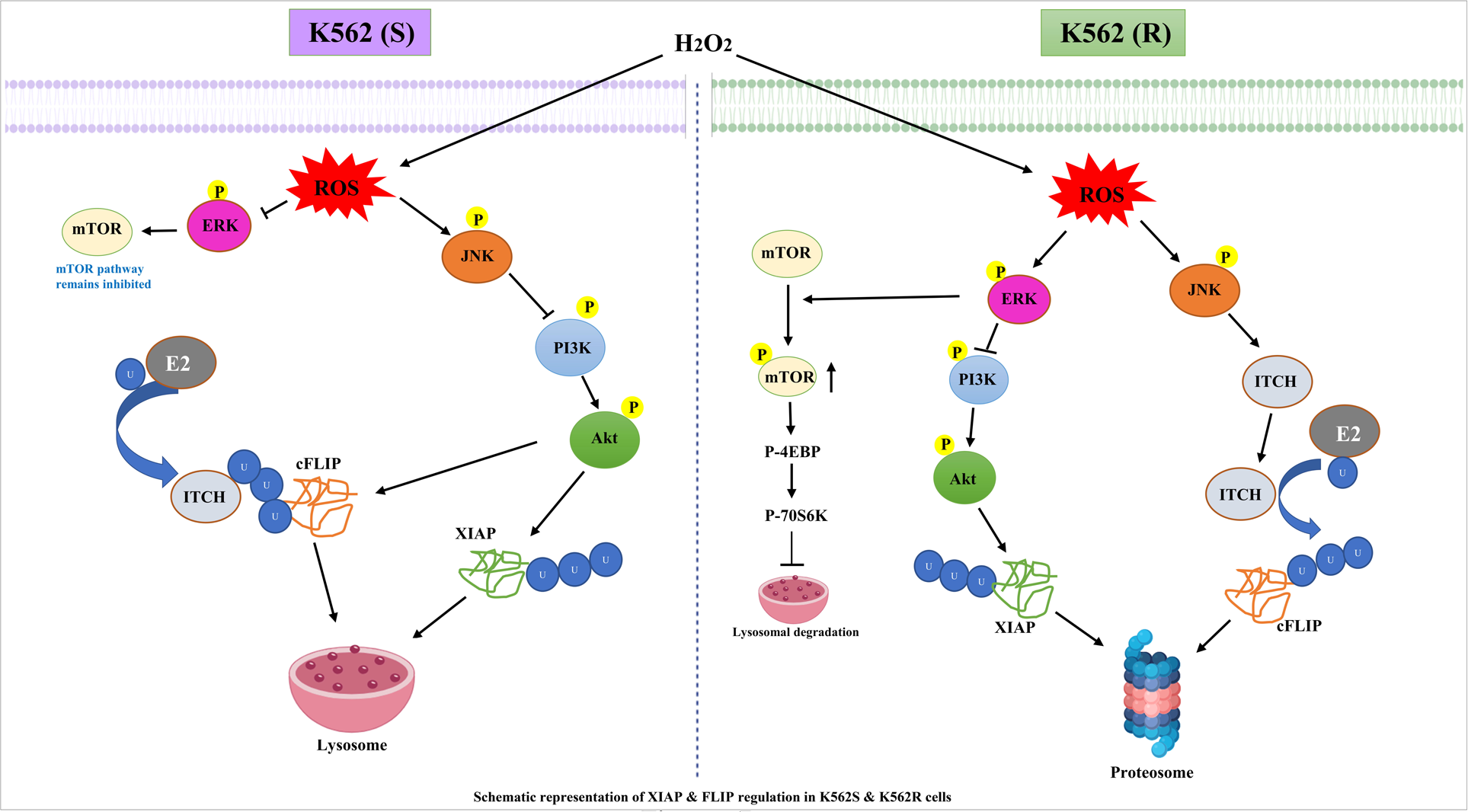
Schematic representation of XIAP and FLIP regulation in K562S & K562R cells. ROS mediates XIAP and FLIP degradation via lysosomal pathway in imatinib sensitive CML cells (K562S) whereas it happens through the proteasomal pathway in imatinib resistant CML cells(K562R) due increased active mTOR pathway which inhibit lysosome-autophagy pathway mediated degradation of FLIP and XIAP in K562R cells.

The final link that we tried to draw is whether this enhanced mTOR activation in K562R cells is responsible behind its Imatinib-resistance. Therefore, we treated both K562S and K562R cells with either 2μM Imatinib alone or in combination with mTOR inhibitor Rapamycin and analyzed the cells annexin V/PI binding assay in flow cytometer for apoptosis. Astonishingly, we have found that K562R cells showed apoptosis markedly like Imatinib sensitive cells (K562S) when rapamycin was pre-treated before imatinib treatment (Fig. 6C). These data conclusively told that mTOR was responsible for Imatinib resistance in K562R cells. Since ROS was showed to be involved in the higher endogenous mTOR phosphorylation, we checked Imatinib resistance after NAC treatment. However, with a lot of surprise, we didn’t find K562 R cells to be converted into sensitive cells like the way rapamycin did (Fig 6 D). Lastly, when cells were pre-treated with ROS scavenger NAC, XIAP and FLIP degradation got reversed (Fig. 6E).

## 5 Discussion

Protein homeostasis (proteostasis) is an important cellular phenomenon in which selective degradation of damaged and aggregated proteins maintain the normal physiological homeostasis. Physiological conditions like developmental changes, adaptation to environmental stress conditions, as well as the accumulation of misfolded or damaged proteins affect the proteostasis. There are two major ways by which protein turnover happens inside the cell. They are the ubiquitin-proteasome system (UPS) and a lysosomal degradation pathway termed macro-autophagy (hereafter autophagy). The small and short-lived proteins take the UPS route (101) and for large proteins as well as aggregates takes the autophagy route.

However, so far, the regulation for targeting selected proteins to any particular degradation route remained largely unknown. Here, in this study, we have addressed this regulation of selection protein degradation pathways. We have selected two important anti-apoptotic proteins XIAP and FLIP as these are the two proteins have been reported by us earlier to be degraded by proteasomal pathway in Imatinib resistant CML cells K562. Now, we took Imatinib resistant K562 (K562R) and Imatinib sensitive K562 (K562S) as two model system for showing the regulation of two different degradation pathways. In our study we found that H_2_O_2_ down regulated both XIAP and FLIP dose and time dependently in K562S and K562R cells, and no significant alteration was observed at transcript level. It was also proved that cells were perfectly viable at the dose of 30 μM H_2_O_2_ thus cell death was not the reason behind this downregulation. However, when we compared the time kinetics of this ROS mediated downregulations, we found difference in the time point at which degradation starts in K562S and K562R cells which indicated that there might be differential regulation or different degradation pathway for these two proteins in K562S and K562R cells. studies with proteasome and lysosome specific inhibitors along with our immunoprecipitation study and western blot analysis with autophagy markers p62 and LC3BII conclusively told that XIAP and FLIP were degraded by H_2_O_2_ via lysosomal pathway in K562S cells and via proteasomal pathway in K562R cells.

As we have discussed earlier that both degradation pathways use ubiquitin to mark the desired proteins for degradation. The ubiquitination of any proteins depends on three enzymes-E1 Ubiquitin activating enzyme, next E2 Ubiquitin conjugating enzyme and E3 Ubiquitin ligase. Among these three, E3 ubiquitin ligase is very much precise to its substrates and thus there are abundant number of ubiquitin ligases to meet this specificity. According to our previous study, E3 ubiquitin ligase ITCH is involved in the ubiquitination of FLIP and XIAP itself act as a ubiquitin ligase and is responsible for its own ubiquitination in K562R cells. In the latter case Akt acts as a stabilizer of XIAP by binding to it[25]. Here also we have found ITCH as ubiquitin ligase for FLIP and XIAP as self-ubiquitin ligase here in K562S. Thus, we have confirmed that altered ubiquitination is not reason behind the different degradation pathway selection in K562R and K562S cells. Therefore, we hypothesize that there must be difference in the signaling pathway which is responsible for these differential control of degradation in K562R and K562S cells.

Since PI3K/Akt and mTOR pathway has been very influential in regulating Ubiquitin proteasome system[26], and autophagy [27], we found it worth checking. We observed that H_2_O_2_ reduced both PI3K and Akt phosphorylation keeping its protein level intact and also found that these dephosphorylation and inactivation is responsible for XIAP and FLIP downregulation because Wortmannin, a well-known inhibitor of PI3K decreased the XIAP and FLIP level either alone or in combination of H_2_O_2_. However, interestingly, this comparison clearly showed that indigenous phosphorylation level of both PI3K and Akt is higher in K562S cells compared to K562R. More interestingly, PI3K and AKT have been found to regulate XIAP in both drug resistant and sensitive cells but only XIAP in K562S cells. Therefore, we concentrated in these pathways to identify the difference in regulation.

One of the most affected pathways by PI3K/Akt is mTOR pathway. when we checked the components of mTOR pathway, we found H_2_O_2_ significantly activated the mTOR pathways in only K562R cells as phosphorylation of mTOR, 4EBP and P70S6K increased with H_2_O_2_ in K562R cells but not in K562S cells. Is this drug-resistant cell-specific mTOR pathway activation responsible for proteasome mediated XIAP and FLIP degradation instead of lysosomal degradation? To answer this, we inhibited mTOR as well as proteasomal degradation pathway with respective inhibitors and found that still H_2_O_2_ was successful in degrading XIAP and FLIP in K562R cells in spite of proteasomal pathways inhibition. Of note, we have already shown that XIAP and FLIP got degraded via proteasomal pathway in K562R cells. We further proved by western blot analysis of autophagy marker p62, LC3BII that autophagy had become activated when both MG132 and Rapamycin were applied in combination during ROS stress. These clearly concluded that mTOR activation leads to suppression of autophagy and as a result, XIAP and FLIP were degraded via lysosomal pathway under ROS stress. This finding also supported previous literatures that soluble substrates are mainly degraded by proteasomes, whereas large insoluble aggregated proteins are generally engulfed by phagophores targeting substrates to the vacuole/lysosome for their destruction[28]. In our case, the XIAP and FLIP degradation which was supposed to take autophagy pathway for destruction, deviated to proteasomal degradation pathway because of activation of autophagy suppression by mTOR.

Next, we tried to find out why mTOR pathway is activated in K562R cells but not in K562S cells. It has been well established by many previous studies, that MAPK pathway component ERK and JNK was downstream to the ROS signaling pathway[29,30]. MAPK pathway consists of three independent components-JNK, ERK and p38 which get activated by phosphorylation. Here, we found that JNK inhibition completely reversed the XIAP and FLIP degradation in K562S cells but inhibition of the other two components failed to do so. On the other hand, JNK inhibition reversed FLIP degradation and ERK inhibition reversed XIAP degradation in K562R cells. Our Knocked down and ectopic overexpression also corroborated our findings that JNK activation is responsible for XIAP and FLIP degradation by H_2_O_2_ in K562S cells.

Here in K562S cells, our data with ectopic overexpression of JNK suggested that JNK activation by H_2_O_2_ led to PI3K and Akt inhibition by decreasing their phosphorylation. Now previous literature suggested that Akt, when phosphorylated bind to XIAP and protect it from degradation[21]. Therefore, decrease in phospho Akt by JNK released XIAP from Akt and activated its self-Ubiquitin ligase activity and increased its lysosomal degradation. On the other hand, JNK increased FLIP degradation by releasing its ubiquitin ligase ITCH from the stronghold of AKT and making them free for its ligation activity.

We also found that H_2_O_2_ dose dependently activated JNK but inhibited ERK in K562S cells whereas it dose-dependently activated both ERK and JNK in K562R cells. Therefore, we clearly found the differential effect of H_2_O_2_ on ERK and similar effect on JNK in K562S and K562R cells. Interestingly, when we knocked down ERK, mTOR activation didn’t happen even after administration of H_2_O_2_ and autophagy was activated more similar in the line of Rapamycin effect. Therefore, it could be concluded here that ERK activation led to mTOR activation and suppression of autophagy in K562R cells. Since ERK got inhibited with H_2_O_2_ in K562S cells, autophagy remained high there and thus the XIAP and FLIP were degraded via autophagy there. Therefore, our data suggested a very interesting role of Erk. Erk had a dual role in K562R cells where it promoted XIAP degradation by inhibiting Akt and also suppressed autophagy by activation of mTOR. However, Erk had no role to play in the degradation of either XIAP or FLIP but its inhibition kept the mTOR suppressed and thus autophagy remained activated.

Since, all of these data so far concluded that H2O2 mediated activation of mTOR in K562R cells is responsible for selection of different degradation pathway of XIAP and FLIP in K562R, we checked the endogenous H2O2 level in these two cells and found that inherent H_2_O_2_ levels were almost double in K562R cells than K562S cells. Moreover, very interestingly, we found that mTOR inhibition by rapamycin resulted in the sensitization of Imatinib-resistant K562R cells to Imatinib. Not only that, we found that ROS scavenger NAC reversed endogenous high mTOR phosphorylation in K562R cells and H2O2 mediated XIAP and FLIP downregulation in K562S cells.

## 6 Conclusion

In conclusions, we have found that ROS mediates XIAP and FLIP degradation via lysosomal pathway in imatinib sensitive CML cells whereas it happens through the proteasomal pathway in imatinib resistant CML cells. The selection of proteasomal pathway for degradation was selected in K562R cells because lysosomal pathway remains inhibited by activated mTOR in K562R cells. Since mTOR remained inhibited in K562S cells, the lysosomal pathway was by default selected for degradation of XIAP and FLIP due to ROS stress. mTOR activation in K562R cells was the result of activated ERK whereas ERK remained inhibited in K562S cells resulting in mTOR inhibition. Quite astonishingly, it has been found that mTOR is not only the deciding factor for degradation pathway selection, mTOR activation also contributed to resistance to Imatinib in K562R cells.

Thus, we showed for the first time that ROS activates differential proteostasis in Imatinib sensitive and Imatinib-resistant CML cells. We showed for the first time that mTOR is a deciding factor for selection of protein degradation pathway. Finally, this work, for the first time, showed that mTOR activation plays a crucial role in the development of resistance to Imatinib.

## Declaration of competing interest

None

## Acknowledgments

We thank Dr. Pralay Majumder of Presidency University for his valuable input in immune fluorescence microscopy. This research work was monetarily supported by the Science and Engineering Research Board, Department of Science and Technology (grant no SB/YS/LS-123/2014 & EEQ12019/000153), Govt. of India; DBT-BUILDER (grant no BT/INF/22/SP45088/2022) program, Department of Biotechnology, Govt. of India. RR is supported by predoctoral fellowship from the Council of Scientific and Industrial Research (New Delhi, India). TP was supported by doctoral fellowships from DBT, India, SS is supported by project fellowship from DST-SERB-EEQ Govt. of India.

---

**Supplementary Figure 1.**
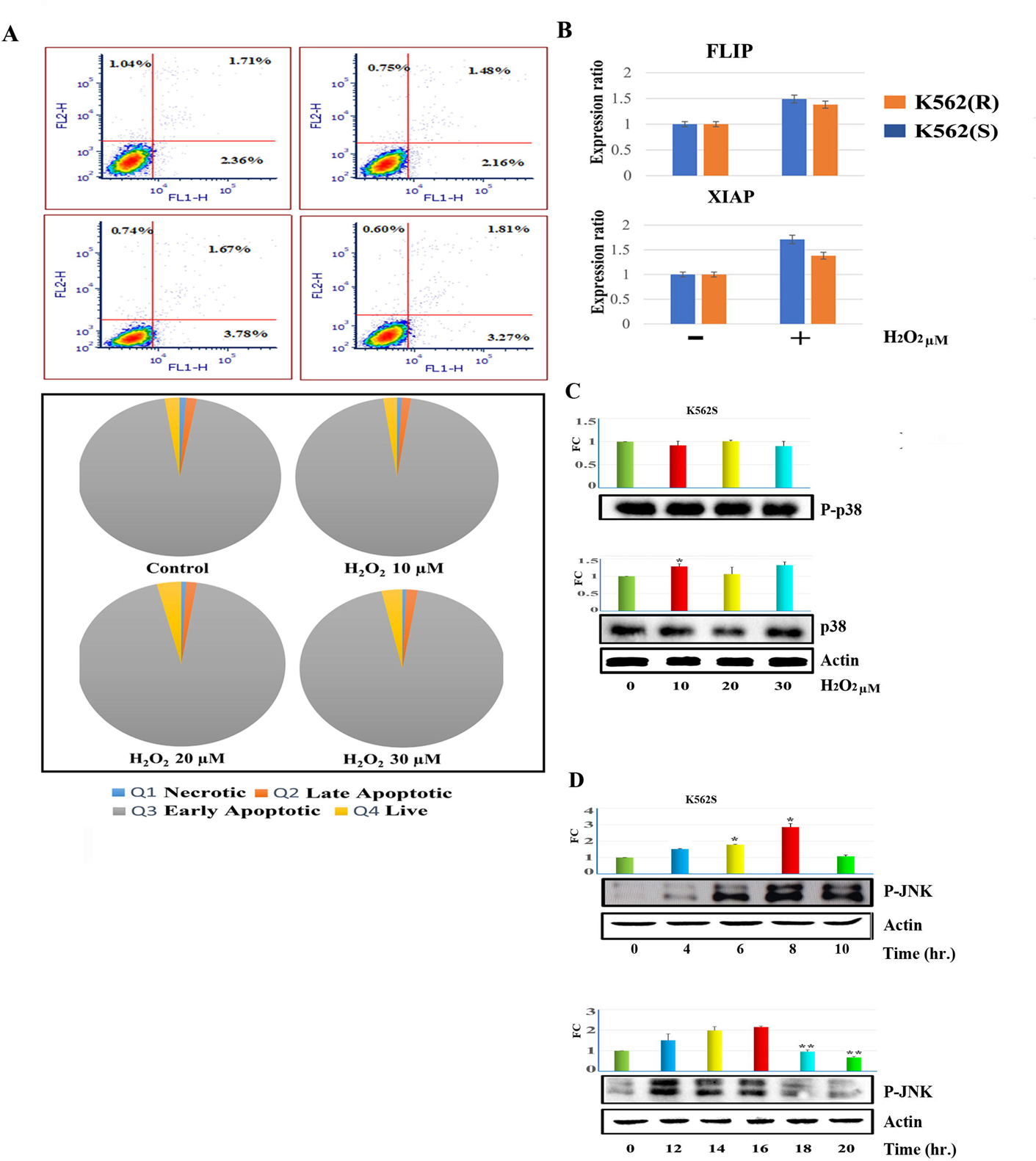
A) Imatinib sensitive K562 cells were treated with specified doses of H_2_O_2_ for 24 hrs, then flow cytometry was performed by subjecting the cells to Annexin V/PI binding assay. The data is representative of three independent experiments. Top panel is the graphical illustration of several population in pie chart and lower panel denotes demonstrative dot plots. B) Both Imatinib sensitive and resistant K562 cells were treated with 30 μM H_2_O_2_ for 14 hours and then total RNA was extracted from cells. Relative levels of endogenous FLIP and XIAP mRNA were evaluated by Real time PCR and graphically denoted as fold change. Data represent mean ± SD of three independent experiments. C) K562(S) cells were incubated with different doses of H_2_O_2_ for 14 hour and whole cell lysate were subjected to western blot analysis with anti P38 and anti p-P38 antibodies. γ Actin was used as loading control. Data were presented graphically as protein level fold change (FC) after densitometry analysis and indicate mean ± SD of three independent experiments. *p < 0.05, **p < 0.01, ***p < 0.001 D) Time kinetics of p-JNK was measured using western blot analysis from imatinib sensitive K562 cells after treating the cells with 30 μM H_2_O_2_ for the specified time points. As loading control γ Actin was used. Data were presented graphically as protein level fold change (FC) after densitometry analysis and indicate mean ± SD of three independent experiments. *p < 0.05, **p < 0.01, ***p < 0.001

## Notes

### Competing Interest Statement

The authors have declared no competing interest.

